# *Atp13a5* Marker Reveals Pericytes of The Central Nervous System in Mice

**DOI:** 10.1101/2021.07.09.451694

**Authors:** Xinying Guo, Tenghuan Ge, Shangzhou Xia, Haijian Wu, Mark Colt, Xiaochun Xie, Bangyan Zhang, Jianxiong Zeng, Jian-Fu Chen, Donghui Zhu, Axel Montagne, Fan Gao, Zhen Zhao

**Affiliations:** Center for Neurodegeneration and Regeneration, Zilkha Neurogenetic Institute and Department of Physiology and Neuroscience, Keck School of Medicine, University of Southern California, Los Angeles, California, 90033, USA; Department of Neurosurgery, Second Affiliated Hospital, School of Medicine, Zhejiang University, Hangzhou, Zhejiang, 310009, China; Neuroscience Graduate Program, Keck School of Medicine, University of Southern California, Los Angeles, California, 90033, USA; The Key Laboratory of Animal Models and Human Disease Mechanisms of Chinese Academy of Sciences & Yunnan Province, Kunming Institute of Zoology, Chinese Academy of Sciences, Kunming, Yunnan 650201, P. R. China; Center for Craniofacial Molecular Biology, University of Southern California, Los Angeles, California, 90033, USA; Department of Biomedical Engineering, Stony Brook University, Stony Brook, NY, United States; Centre for Clinical Brain Sciences, UK Dementia Research Institute at University of Edinburgh, Edinburgh, United Kingdom; Caltech Bioinformatics Resource Center, California Institute of Technology, Pasadena, CA 91125

## Abstract

Perivascular mural cells including vascular smooth cells (VSMCs) and pericytes are integral components of the vascular system. In the central nervous system (CNS), pericytes are also known as the guardian of the blood-brain barrier, blood-spinal cord barrier and blood-retinal barrier, and play key roles in maintaining cerebrovascular and neuronal functions. However, the functional difference between CNS and peripheral pericytes has not been resolved at the genetic and molecular levels. Hence, the generation of reliable CNS pericyte-specific models and genetic tools remains very challenging. Here, we report a new CNS pericyte marker in mice. This cation-transporting ATPase 13A5 (*Atp13a5*) marker is highly specific to the pericytes in brain, spinal cord and retina. We generated a transgenic model with a knock-in tdTomato reporter and Cre recombinase. The tdTomato reporter reliably labels the CNS pericytes, but not found in any other CNS cell types including closely related VSMCs, or in peripheral organs. More importantly, *Atp13a5* is turned on at embryonic day E15, suggesting brain pericytes are shaped by the developing neural environment. We hope that the new tools will allow us to further explore the heterogeneity of pericytes and achieve a better understanding of CNS pericytes in health and diseases.

## INTRODUCTION

Pericytes are vascular mural cells that play key roles in vascular development and the maintenance of microvascular functions (Armulik et al., 2011; Sweeney et al., 2016). They cover microvessels including arterioles, venules and capillaries, while vascular smooth muscle cells (VSMCs) occupy large-diameter arteries and veins (Sweeney et al., 2016). In the central nervous system (CNS), pericytes are vital integrators of neurovascular functions (Hartmann et al., 2021; Nikolakopoulou et al., 2019), and indispensable for a functional blood-brain barrier (BBB) (Armulik et al., 2010; Bell et al., 2010). It is proposed based on early genetic lineage tracing studies that brain pericytes may originate from neural crest cells, while peripheral ones mainly arise from mesothelium (Armulik et al., 2011; Yamazaki and Mukouyama, 2018). However, no genetic marker has been identified so far for a clear classification between CNS and pericyte pericytes, which also become a major hurdle for genetic manipulations and linage tracing of CNS pericytes (Sweeney et al., 2016).

Various genetic markers of pericytes have been tested in the past decade. Platelet-derived growth factor receptor beta (PDGFRβ) is perhaps the most well-known molecular marker for pericytes, as PDGF-B/PDGFRβ signaling is essential for its fate determination (Armulik et al., 2010; Bell et al., 2010). In addition, chondroitin sulfate proteoglycan 4 (*CSPG4*), desmin, vimentin, regulator of G protein signaling 5 (*RGS5*), CD13/aminopeptidase N (*APN*) are used as well. However, almost all these markers are expressed in VSMCs and/or fibroblasts. Therefore, transgenic mouse models based on these alleles, including *Cspg4-Cre* and *Cspg4-dsRed* (Zhu et al., 2008), *Pdgfrb-EGFP and Pdgfrb-Cre* (Gerl et al., 2015)*, Rgs5-EGFP* (Nisancioglu et al., 2008), all have limitations when applied to brain pericytes. In addition, these markers can hardly differentiate CNS pericytes from peripheral ones, let alone teasing out the inherent heterogeneity (Vanlandewijck et al., 2018). Recently, we reported using a double-promoter approach with both the *Pdgfrb* and *Cspg4* promoters to control Cre recombinase expression (Nikolakopoulou et al., 2019). This model exhibits restricted Cre activity in pericytes compared to VSMCs, but leakage in peripheral organs remains (Nikolakopoulou et al., 2019), and the sophisticated design with two promoters and two recombinases (Cre and Flp) also limits its applications.

To address this gap, we compiled multiple mouse transcriptomic datasets, and identified that *Atp13a5* is much more specific to brain pericytes than other current markers. *Atp13a5* encodes a member of the P5 subfamily of P-type ATPases, and is predicted to be a cation transporter (Sørensen et al., 2010). *Atp13a5* is also developmentally regulated, and its appearance coincides with BBB establishment around embryonic day E15. Next, we generated an *Atp13a5-2A-CreERT2-IRES-tdTomato* knock-in model, by replacing the endogenous stop codon with a cassette with a self-cleaving 2A peptide sequence and an internal ribosome entry site (IRES) for cistronic expression of both Cre recombinase and tdTomato reporter. Profiles of the tdTomato reporter in this new model are on a par with the bioinformatic results, as they are only found in the CNS and colocalized exclusively with CD13^+^ pericyte profiles. More importantly, *Atp13a5-driven* tdTomato reporter and Cre-dependent GCaMP6 reporter are completely absent in peripheral organs, suggesting that this genetic tool based on *Atp13a5* is successful and can be utilized for genetic manipulations and linage tracing of CNS pericytes *in vivo*.

## Results

### *Atp13a5* is specifically expressed in mouse brain pericytes

To identify a new marker for mouse brain pericytes, we compared five different transcriptomic datasets (Armulik et al., 2010; Daneman, 2010; He et al., 2016; Song et al., 2020; Ximerakis et al., 2019), and found that only 16 genes were commonly identified among these studies (**Sup. Fig. 1A** and **Sup. Table1**). More importantly, we analyzed the single cell RNA sequencing (scRNA-seq) data of brain vasculature (Vanlandewijck et al., 2018). 3,186 single-cell transcriptomes were collected for the secondary analysis using a new R Seurat Package (Butler et al., 2018; Stuart et al., 2019). In the Uniform Manifold Approximation and Projection (UMAP), 9 major cell types were separated into clusters, which including pericytes (PC), vascular smooth muscle cells (VSMC), arterial endothelial cells (aEC), venous endothelial cells (vEC), capillary endothelial cells (capEC), Oligodendrocytes (Oligo), fibroblast (FB), microglia (MG), astrocytes (AC) (**Fig. 1A**), based on specific genetic markers (**Sup. Fig. 1B** and **Sup. Table2**). The top 3 gene markers in each cell type showed their distinct cluster-specific expression patterns (**Fig. 1B**). For example, vitronectin (*Vtn*) and the inwardly-rectifying potassium channel *Kcnj8* are pan-pericyte markers, enriched in both brain pericytes and lung pericytes; however, they both express in a subset of VSMC as well (**Fig. 1C**).

**Figure 1.**
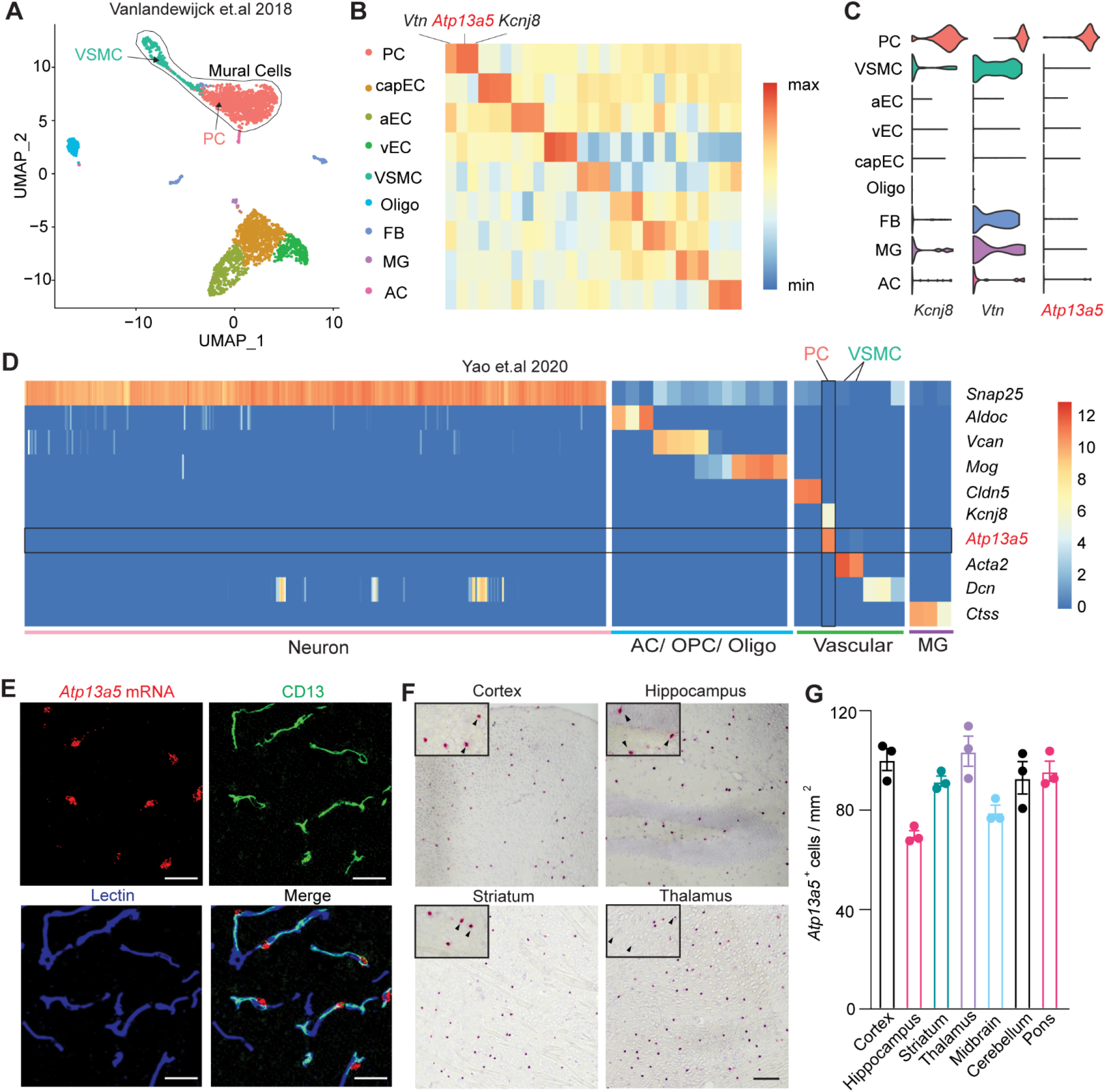
*Atp13a5* is specifically expressed by brain pericytes. (**A**) UMAP of brain vasculature transcriptomes. Mural cells are marked by black line. (**B**) Gene-expression heatmap of the top 3 markers genes in each cluster. (**C**) Violin plots showing the distribution of expression level of the top 3 pericyte markers across all 9 cell types. (**D**) Gene expression heatmap of representative genes in Allen Institute’s dataset. PC: Pericytes; capEC: capillary endothelial cells; aEC: arterial endothelial cells; vEC: venous endothelial cells; VSMC: vascular smooth muscle cells; Oligo: Oligodendrocytes; FB: Fibroblast; MG: microglia; AC: astrocytes; OPC: Oligodendrocyte progenitor cells. (**E**) Representative images for *Atp13a5* mRNA expression (Red) and immunostaining for CD13^+^ pericytes (Green) and Lectin^+^ endothelia cells (Blue) in cortex. Scale bar: 50 μm. Sections: 15 μm thickness. (**F**) Representative images for *Atp13a5* mRNA expression (Red) in various mouse brain regions. Scale bar: 100 μm. Sections: 10 μm thick. (**G**) Number of *Atp13a5*+ cells per mm^2^ in different mouse brain regions. n = 3 mice. Data are presented in mean ± SEM.

Interestingly, we found *Atp13a5*, which encodes a member of the P-type ATPases, is highly expressed in brain pericytes but not in other vasculature cells including VSMCs (**Fig. 1C**). In addition, we examined a dataset of 1,093,785 cells from multiple cortical and hippocampal areas (Yao et al., 2020) and found that *Atp13a5* is indeed specific to brain pericytes (**Fig. 1D**). To further validate its transcripts in brain, we used fluorescent *in situ* hybridization (FISH) with RNAscope probes and found that *Atp13a5* mRNA is colocalized exclusively with CD13^+^ pericyte profiles in the cortex, but not with other cells including endothelial cells (**Fig. 1E**). RNAscope results also showed that *Atp13a5-positive* cells are detected throughout the brain, including cortex, hippocampus, striatum, thalamus, midbrain, pons and cerebellum (**Fig. 1F-G** and **Sup. Fig. 1C**). These results match with the *Atp13a5 in situ* hybridization data from the Allen Brain Atlas (**Sup. Fig. 1D** and **E**).

### Atp13a5 expression is highly enriched in the mouse brain

To compare *Atp13a5* expression between mouse brain and peripheral organs, we analyzed the Tabula Muris dataset (The Tabula Muris Consortium et al., 2018). After visualizing all cells with UMAP and grouped them with unbiased, graph-based clustering (**Fig. 2A-B**), we found that *Atp13a5-*expressing cells are only present in the pericyte clusters marked by *Rgs5, Kcnj8*, and *Pdgfrb* (**Fig. 2C** and **Sup. Fig. 2A-B**). More importantly, *Atp13a5-*expressing cells are mainly of brain origin (**Fig. 2D**), whereas other pericyte markers including *Rgs5, Kcnj8*, and *Pdgfrb* are highly expressed in the periphery tissues as well (**Sup. Fig. 2B-D**). This result is consistent with tissue specific RNAseq data on NCBI (**Sup. Fig. 2E**), indicating that *Atp13a5* is very likely a brain pericyte-specific gene. We further confirmed with FISH with RNAscope probes that *Atp13a5* is highly expressed in brain pericytes, but not found in kidney, liver, and the ventricular wall of heart (**Fig. 2E**).

**Figure 2.**
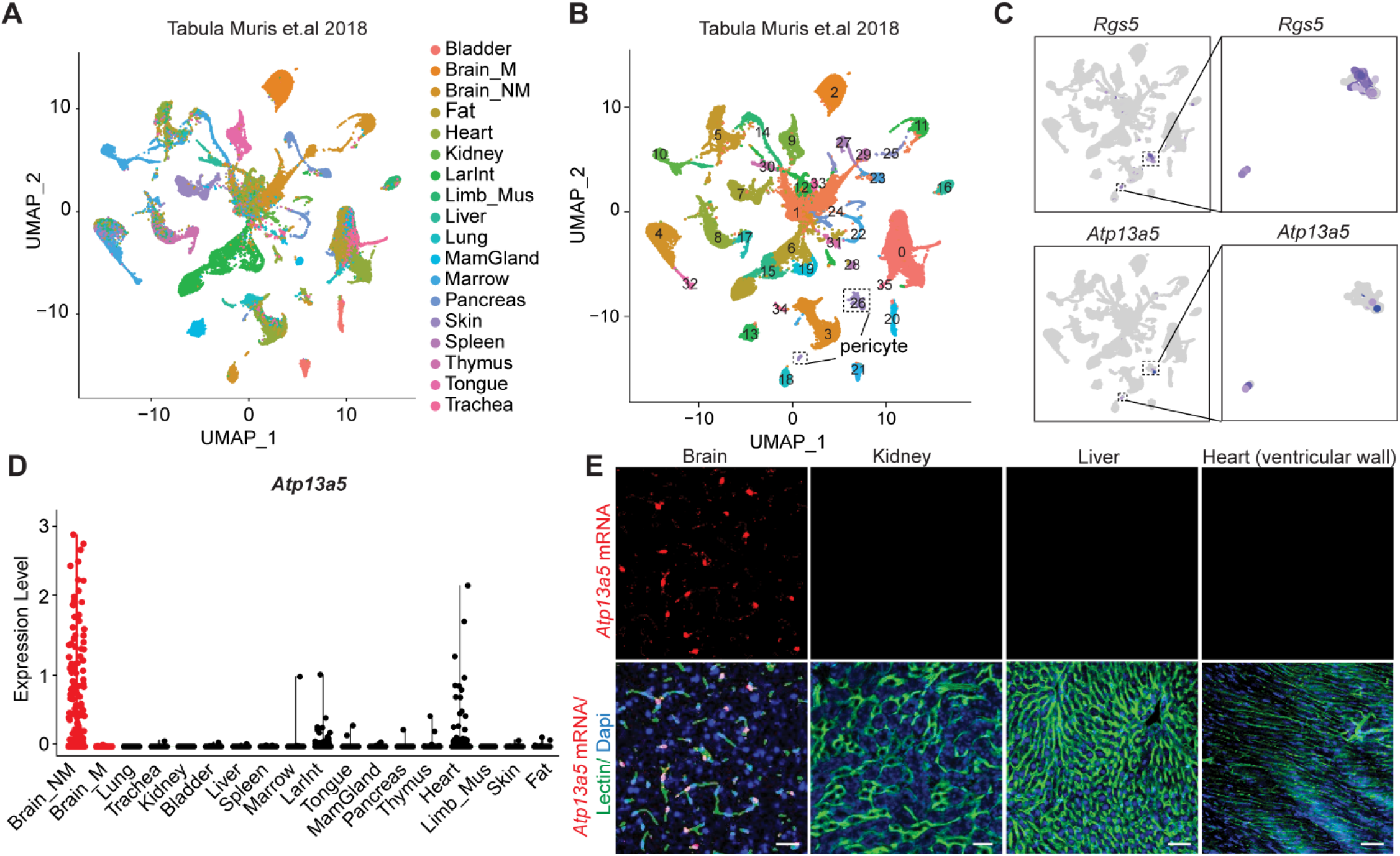
*Atp13a5* is highly enriched in the central nervous system. (**A**) UMAP of all cells from 18 tissues. Each dot was color-coded and annotated by organs. (**B**) UMAP of 53,760 cells. Each dot was color-coded and annotated by graph-based clusters. Pericytes are marked by black box. **(C)**UMAP plots showing scRNA-seq data, colored by gene expression value, showing *Rgs5* and *Atp13a5* expression. (**D**) Violin plots showing *Atp13a5* that distinguish across and within 18 tissues. Red dot indicated the brain tissue. Brain_M: Brain myeloid; Brain_NM: brain non-myeloid; LarInt: large intestine; Mus: muscle; MamGland: Mammary Gland. **(E)**FISH for *Atp13a5* mRNA (Red) and immunostaining for Lectin^+^ endothelial cells (Green) and Dapi (Blue) in the cortex of brain, kidney, liver and the ventricular wall of heart. Scale bar: 50 μm. Sections: 15 μm.

### Generation of *Atp13a5-2A-CreERT2-IRES-tdTomato* knock-in model

Next, we generated a new transgenic model targeting the endogenous *Atp13a5* allele, to carry both Cre recombinase for genetic manipulation and a fluorescence reporter for imaging and lineage tracing (Sjulson et al., 2016). More specifically, this *Atp13a5-2A-CreERT2-IRES-tdTomato* knock-in model harnesses the endogenous *Atp13a5* locus to drive the expression of both Cre and tdTomato, while preserving endogenous *Atp13a5* integrity by using the self-cleaving 2A peptide sequence (Tang et al., 2009) and an internal ribosome entry site (IRES) (Hellen, 2001) (**Fig. 3A**, also see Methods). One F0 founder was selected based on germline transmission and genome sequencing (**Sup. Fig. 3A**); and the F1 generation was further tested with southern blot analysis for the integrity of the knock-in allele (**Sup. Fig. 3B**). Homozygous *Atp13a5-2A-CreERT2-IRES-tdTomato (Atp^tdTtdT^*) mice are viable and appear normal. The knock-in cassette does not affect endogenous *Atp13a5* expression, as validated by quantitative real-time PCR (**Sup. Fig. 3C-D**).

**Figure 3.**
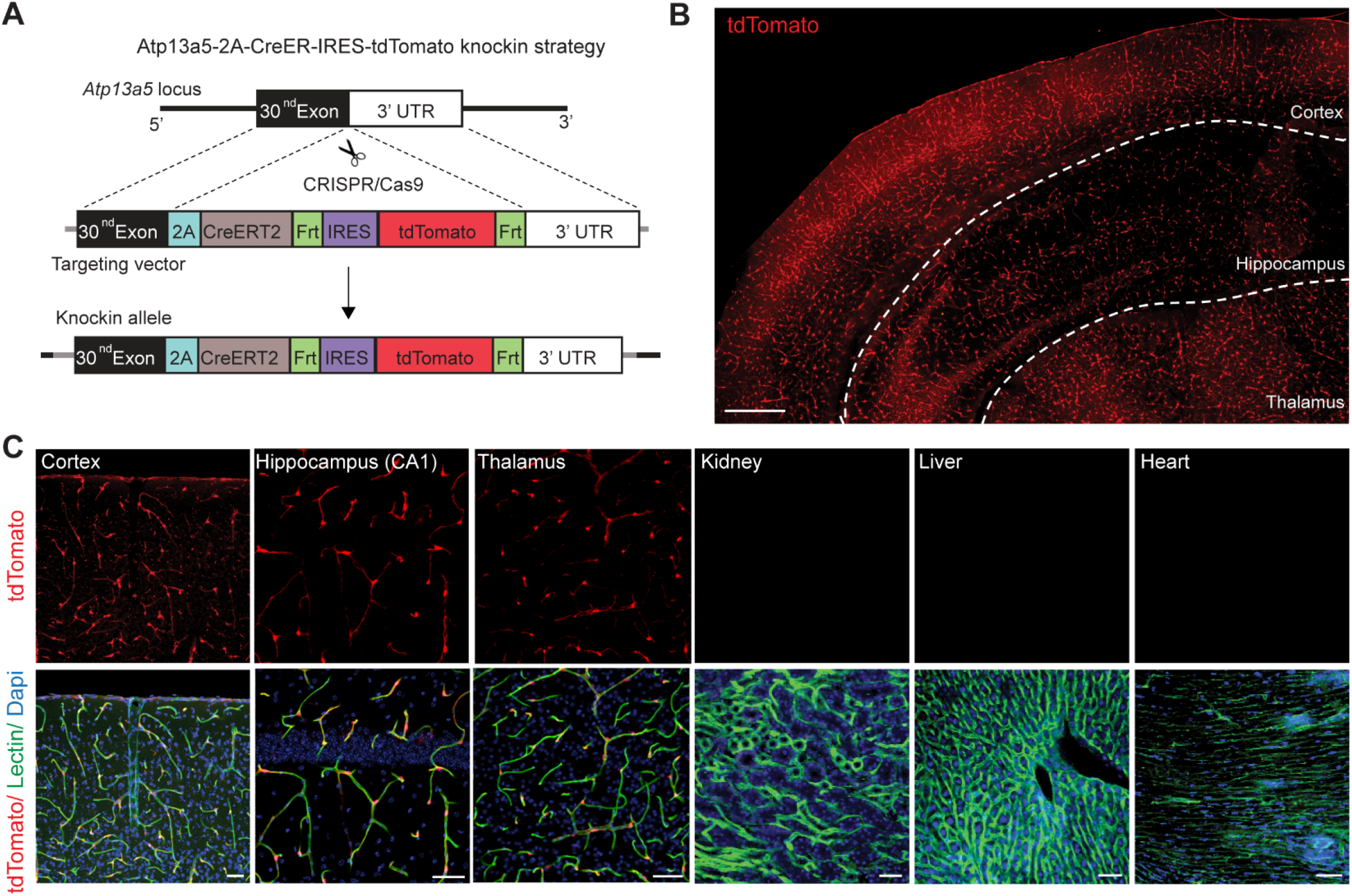
Generation and validation of the *Atp13a5-2A-CreERT2-IRES-tdTomato* model. **(A)** Schematic diagram showing the strategy for generating the *Atp13a5-2A-CreERT2-IRES-tdTomato* knock-in mice. See Methods for more details. **(B)** A representative tiled image of brain section from a heterozygous *Atp13a5-2A-CreERT2-IRES-tdTomato* mouse. Scale bar: 500 μm. **(C**) Representative confocal images of tdTomato, endothelial marker Lectin and Dapi in different tissues from a homozygous *Atp13a5-2A-CreERT2-IRES-tdTomato* mouse, including cortex, CA1 region of hippocampus, thalamus, kidney, liver and heart (ventricular wall). Scale bar: 50 μm.

### *Atp13a5-tdTomato* expressing cells are CNS pericytes

We found that *Atp13a5*-driven tdTomato is reliably expressed in adult heterozygous *Atp^tdT/+^* (**Fig. 3B**) and homozygous *Atp^tdT/tdT^* mice (**Fig. 3C**). The tdTomato profiles cover the Lectin-labelled microvessels throughout the brain regions including cortex, hippocampus and thalamus, but are not seen in peripheral tissues such as kidney, liver, or heart (**Fig. 3C**). Additional immunostaining further confirmed that tdTomato is not expressed in any VCAM1-possitive venules (**Fig. 4A-B**), or smooth muscle cell actin (SMA)-positive arterioles (**Fig. 4C-D**), but only in CD13-positive pericytes in brain (**Fig. 4E-F**). No leakage of reporter expression was found in oligodendrocytes, neurons, microglia or astrocytes (**Sup. Fig. 4A-F**). When compared to the well-characterized *Pdgfrb-EGFP* model (**Sup. Fig. 5A**), the new *Atp13a5-2A-CreERT2-IRES-tdTomato* model has much improved specificity to the brain pericytes, over the peripheral ones (**Sup. Fig. 5B**).

**Figure 4.**
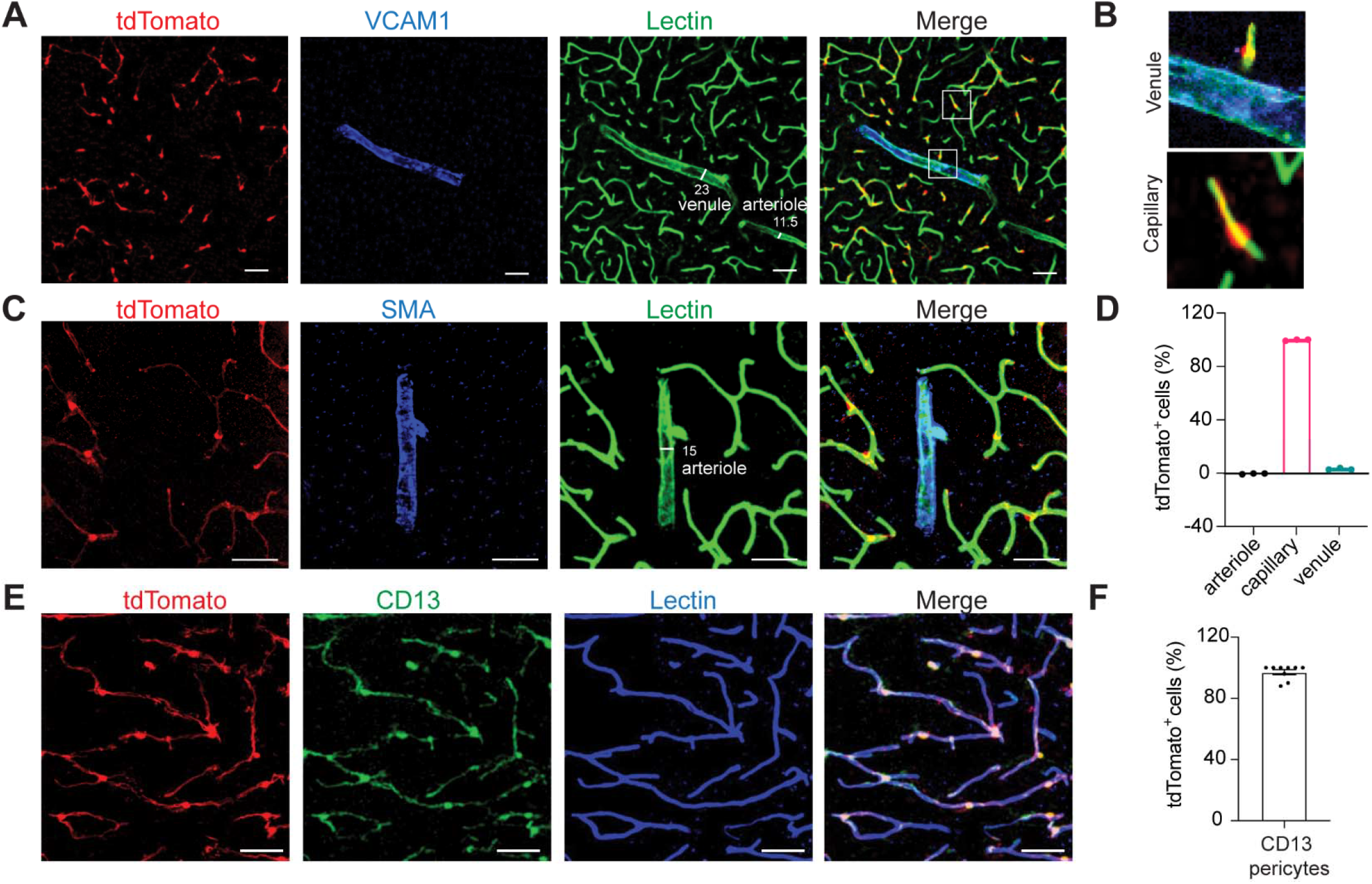
*Atp13a5-driven tdTomato* reporter expression in brain pericytes. **(A)** tdTomato expression on brain capillary of *Atp13a5-2A-CreERT2-IRES-tdTomato* knock-in mice, but not on VCAM1^+^ venules. Scale bar: 50 μm. **(B)** Representative images from the boxed regions in **A**. **(C)** tdTomato expression on brain capillary, but not on SMA^+^ arterioles. Scale bar: 50 μm. **(D)** The percentage of tdTomato^+^ cells distributed among arterioles, capillaries and venules in the cortex. Arteries and arterioles are identified by vessel diameter in combination with the presence of SMA. Veins and venules are identified by vessel diameter in combination with the presence of VCAM1 and the absence of SMA. Lectin^+^ vessels with diameters smaller than 6 μm are considered as capillaries. n = 3 mice **(E)**Colocalization of tdTomato with pericyte marker CD13 (green) on Lectin (blue) positive endothelium. Scale bar: 50 μm. **(F)**Quantification of the percentage of tdTomato^+^ cells in CD13^+^ pericytes. n = 9 mice. Data are presented in mean ± SEM.

To verify the expression of *Atp13a5* marker throughout the CNS, we also examined the spinal cord and retina in *Atp13a5-2A-CreERT2-IRES-tdTomato* model. We found robust tdTomato signals in both white matter and gray matters of the spinal cord (**Fig. 5A-B**), with higher coverage in gray matter. In retina, pericytes expressing *Atp13a5*-tdTomato are found in most of the microvessels (**Fig. 5C-D**), particularly in the ganglion cell layer and inner nuclear layer (**Fig. 5E**). Hence, *Atp13a5*-expressing cells are truly mouse CNS pericytes.

**Figure 5.**
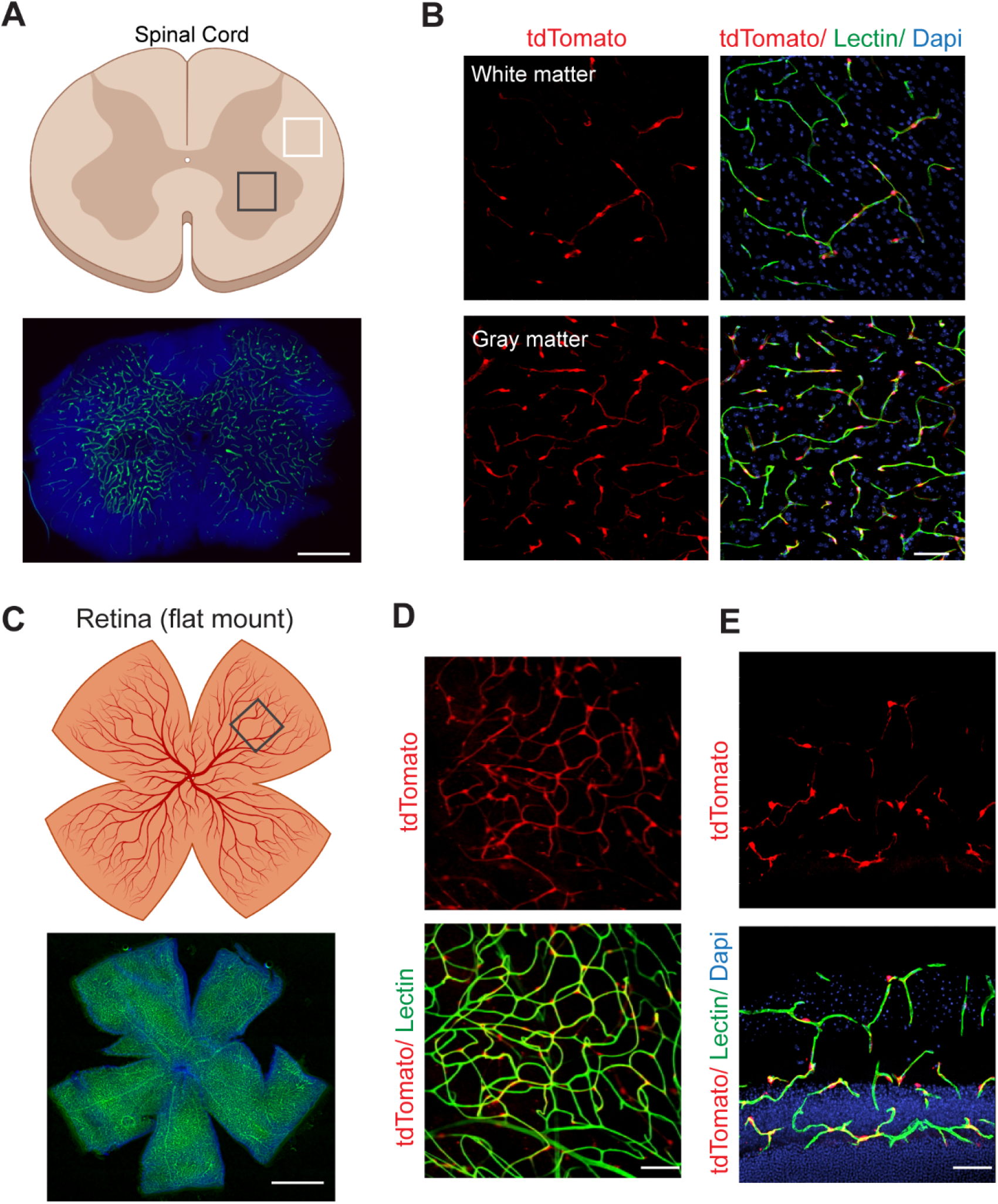
*Atp13a5-driven tdTomato* reporter expression in spinal cord and retina. **(A)** A diagram showing a cross section of mouse spinal cord on top, and lectin angiogram (green) on bottom. Dapi, nuclear staining. Scale bar: 500 μm. **(B)** tdTomato reporter expression in the spinal cord pericytes of *Atp13a5-2A-CreERT2-IRES-tdTomato* knock-in mice. Scale bar: 50 μm. **(C)** A diagram showing a flat mount preparation of mouse retina on top, and lectin angiogram (green) on bottom. Dapi, nuclear staining. Scale bar: 1 mm. **(D-E)** tdTomato reporter expression in the retinal pericytes of *Atp13a5-2A-CreERT2-IRES-tdTomato* knock-in mice. Scale bar: 50 μm. **D**, flat mount; **E**, cross section.

### Developmental regulation of *Atp13a5* marker

Next, we analyzed 24,185 single cells from mouse dentate gyrus (Hochgerner et al., 2018), spanning perinatal (E16.5-P5), juvenile (P18-P23) and adult (P120-P132) ages (**Fig. 6A**). Again, *Atp13a5-* expressing cells are only present in the pericyte cluster (**Fig. 6B** and **Sup. Fig. 6A**). Interestingly, *Atp13a5* expression increased after birth and is sustained throughout adulthood (**Fig. 6C**). We further performed FISH at different ages. By quantifying the density of *Atp13a5* transcripts in CD13^+^ pericytes, we found that *Atp13a5* is expressed in *10% CD13^+^ brain pericytes at embryonic day E15 (**Fig. 6D**); while at birth (P0), most brain pericytes expressed *Atp13a5* (**Fig. 6E**). This period is also critical for BBB establishment (Zhao et al., 2015), suggesting a neuronal environment may contribute to the specialization of brain pericytes. After birth, *Atp13a5* levels continue to increase in brain pericytes until P10; and this level is sustained even after 18 months (**Fig. 6D-E**). The developmental regulation of *Atp13a5* is also recapitulated by the transgenic reporter. tdTomato^+^ brain pericytes are not seen at E12 (**Fig. 6F-G**), barely visible at E15 (**Sup. Fig. 6B-C**), and become prominent at E16 (**Fig. 6H-J**). At P0, the tdTomato profiles are seen nearly on all vessels (**Sup. Fig. 6D-E**). Therefore, *Atp13a5* expression is developmentally regulated, but it is a reliable marker for studying pericytes in postnatal, juvenile, adult and aging mice.

**Figure 6.**
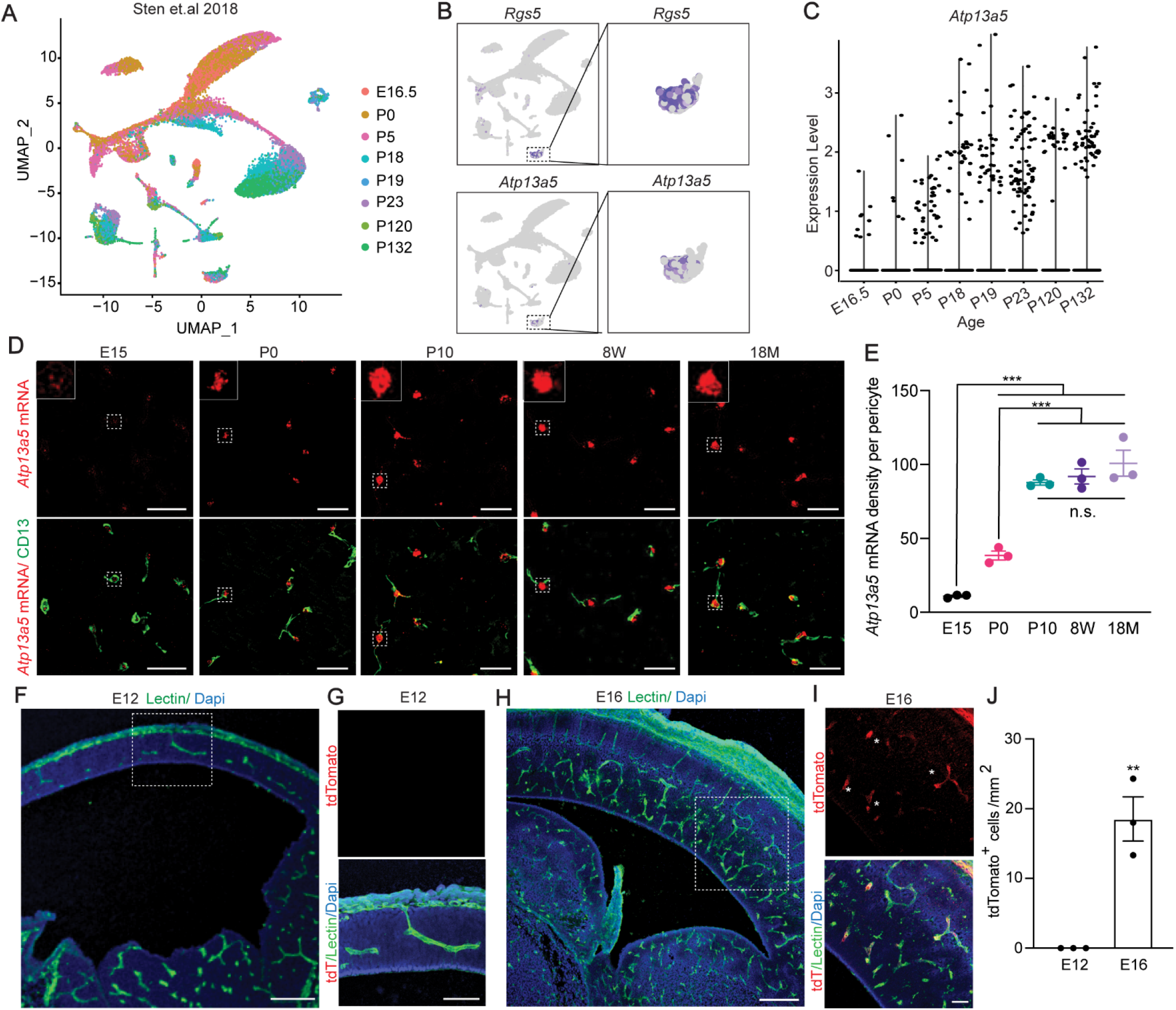
Developmental regulation of *Atp13a5* marker. **(A)** UMAP of 24,185 mouse brain cells from multiple ages. Each dot was color-coded and annotated by different ages. **(B)** UMAP plots showing scRNA-seq data, colored by gene expression value, showing *Rgs5*and *Atp13a5* expression. (**C**) Violin plots showing *Atp13a5* expression in different ages**. (D)** Representative images for *Atp13a5* mRNA expression and CD13 in brain sections from different ages. Scale bar: 50 μm. Sections: 15 μm thick. **(E)** *Atp13a5* expression density per pericyte in various developmental stages. Data are presented in mean ± SEM; n = 3 mice per age group; ***, P < 0.001, n.s., no significant difference, by One-way ANOVA with Tukey Test. **(F)** A representative image of an *Atp13a5-2A-CreERT2-IRES-tdTomato* embryonic cortice at E12. Scale bar: 200 μm. Sections: 30 μm thick. **(G)** High magnification of boxed region in **F**. tdTomato was not detected in cortex at E12. Scale bar: 100 μm. **(H)** A representative image of an *Atp13a5-2A-CreERT2-IRES-tdTomato* embryonic cortice at E16. Scale bar: 200 μm. Sections: 30 μm thick. **(I)** High magnification of boxed region in **H**. tdTomato can be detected in cortex at E16. Scale bar: 50 μm. Sections: 30 μm thick. **(J)** Quantification of tdTomato^+^ pericyte numbers per mm^2^ in E12 and E16 mouse brain. n = 3 mice. **, P < 0.01 by Student’s t-test. Data are presented in mean ± SEM.

### Characterization of the *Atp13a5-CreER* recombinase activity

To test the CreER activity in this *Atp13a5-2A-CreERT2-IRES-tdTomato* model, we crossed it with the Cre-dependent and Tet-controllable Ai162 line with fluorescent calcium-indicator GCaMP6s (Daigle et al., 2018), and induced the CreER activity with tamoxifen administration (**Fig. 7A**, also see Methods). With 4 injections of Tamoxifen, we observed nearly 40% of tdTomato^+^ brain pericytes expressing robust GCaMP6s protein (**Fig. 7B**), but not in peripheral tissues such as heart, kidney or liver (**Fig. 7C-D**). This sparse labeling also allows us to clearly resolve the morphology of single pericytes. While the majority of the tdTomato^+^ and GCaMP6s^+^ double positive profiles exhibit elongated processes covering the microvessels (Type I)(Nikolakopoulou et al., 2019), we also observed pericytes with shorter processes wrapping around the whole microvessel (Type II), as well as a hybrid type with both elongated and wrapping processes (Type III). The type I and type II are known as thin-strand and mesh pericytes, respectively (Berthiaume et al., 2018); however the hybrid type has not been reported before, suggesting the heterogeneity of CNS pericytes could be more complicated than our current understanding. Taken together, our data demonstrated that the new *Atp13a5* marker reveals the CNS pericytes, and *Atp13a5-2A-CreERT2-IRES-tdTomato* model will help us to explore the biology of CNS pericytes *in vivo*.

**Figure 7.**
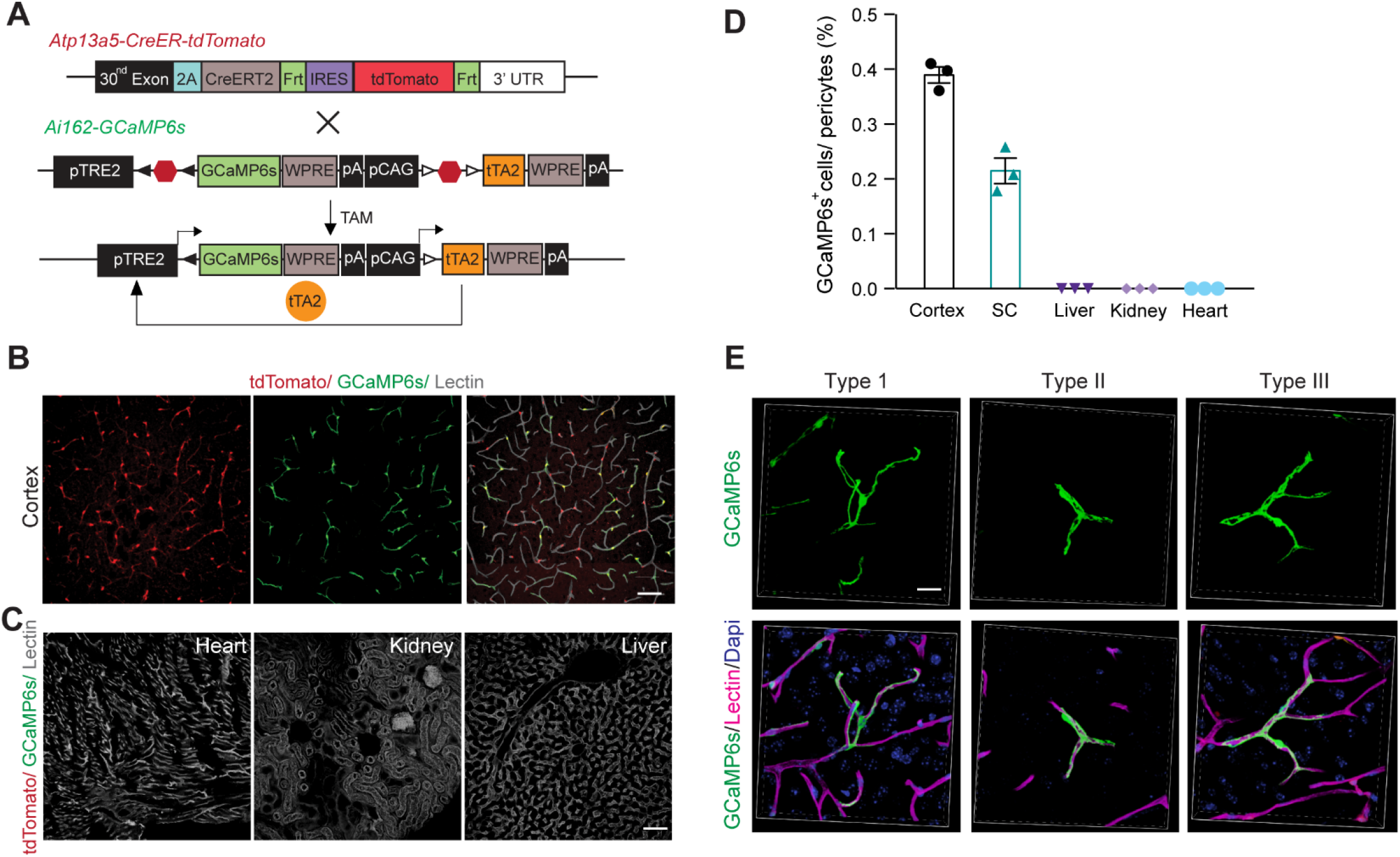
Characterizing the CreER activity. **(A)** Schematic diagram showing the breeding strategy for generating *Atp13a5-2A-CreERT2-IRES-tdTomato; Ail62* mice. **(B)** Representative confocal images of cortical section from a *Atp13a5-2A-CreERT2-IRES-tdTomato; Ai162* mouse, with *Atp13a5*-driven tdTomato (red), tamoxifen-induced GCaMP6s (green) and lectin-labelled endothelial profiles (gray). Scale bar: 50 μm. **(C**) Representative confocal images of heart, kidney and liver sections from a *Atp13a5-2A-CreERT2-IRES-tdTomato; Ai162* mouse, showing no tdTomato or GCaMP6s expression. Lectin (gray): endothelial profiles. Scale bar: 50 μm. **(D**) Quantification of the percentages of tdTomato^+^ and GCaMP6s^+^ double positive cells in pericytes in different organs as indicated. n = 3 mice. Data are presented in mean ± SEM. **(E**) High resolution 3D reconstruction of sparse labelled tdTomato^+^ and GCaMP6s^+^ double positive brain pericytes. Type I: thin-strand pericytes; type II: mesh pericytes; type III: hybrid pericytes. Scale bar: 20 μm.

## DISCUSSION

Brain pericytes regulate vascular development and microvascular functions; however, their identification still requires a combination of inadequate markers shared with other cell types, including closely related fibroblasts and VSMCs. This has become a major hurdle to a better understanding of pericyte biology, heterogeneity and contributions towards human diseases. Here, we identified a novel genetic marker *Atp13a5* for mouse brain pericytes through 8 transcriptomic datasets, and generated a new transgenic tool based on this allele. This *Atp13a5* genetic marker, as well as the *Atp13a5-2A-CreERT2-IRES-tdTomato* knock-in reporter model, demonstrated high specificity to brain pericytes. First, *Atp13a5* is not expressed in VSMCs or perivascular fibroblasts; second, it exclusively labels brain pericytes, but not the peripheral ones. More importantly, when tracing the *Atp13a5*-expressing cells in the reporter model, we surprisingly found that they are exclusively located in the CNS regions including brain, spinal cord and retina; while the pericytes in the peripheral tissues do not express this marker, e.g., heart, kidney and liver. Therefore, we hope that the *Atp13a5* marker and the *Atp13a5-2A-CreERT2-IRES-tdTomato* knock-in model will become useful tools for pericyte research.

The mammalian BBB is central to the CNS health and functions, which is established during embryonic development and influenced by the neural environment (Zhao et al., 2015). The developmental origin of pericytes seems to be rather complex (Yamazaki and Mukouyama, 2018); and their heterogeneity has only been confirmed between central and peripheral tissues, but not within the CNS (Vanlandewijck et al., 2018). The finding of *Atp13a5-based* brain pericyte heterogeneity now allows us to examine brain pericytes development and the fate determination of BBB pericytes more accurately, at anatomical and transcriptional levels. More specifically, we found *Atp13a5* is turned on around embryonic day E15 in mice, when the BBB starts to be functional, which is different from other known markers (Jung et al., 2018). This suggests that brain pericytes can also be influenced by the local neural environment and metabolic reprogramming (Sheikh et al., 2020), and switch their property towards more specialized brain pericytes. They could be regulated by the Wnt signaling and induced by the neuronal activity, just like the BBB endothelium (Zhao et al., 2015), or by completely new mechanism yet to be deciphered. Nevertheless, more comprehensive decoding of the transcriptional differences between the CNS and peripheral pericytes remains to be explored. Perhaps, a combined model with *Atp13a5^tdT^* and *Pdgfrb*-EGFP might help, particularly at the embryonic stages when developing brain pericytes are quite heterogenous (Fig. 6).

Although pericytes and VSMCs are anatomically positioned at different locations of the vascular segment and are functionally different, their biological identifies are similar in many ways. This has made it difficult to understand the *in vivo* functions of brain pericytes and perhaps led to certain disagreements regarding the role of pericytes in regulating cerebral blood flow (Hall et al., 2014; Hartmann et al., 2021; Hill et al., 2015). This issue perhaps can be resolved now with the new *Atp13a5-2A-CreERT2-IRES-tdTomato* knock-in model. It is a versatile tool with not only an endogenous tdTomato reporter for tracing CNS pericytes, as well as a tamoxifen-inducible Cre recombinase for effective genetic manipulation, specifically in these cells. The robustness of the tdTomato signal in this model enables cell sorting for molecular profiling and cell culture without the need for antibodies, as well as intravital imaging and lineage tracing *in vivo*. More importantly, the CreERT2 element will provide us an inducible approach *in vivo*, including genetic manipulation and ablation, especially in cases when brain VSMCs or peripheral pericytes may confound interpretations. In addition, the IRES-tdTomato sequence is further flanked by two flp recombinase target (frt) sites for future removal of the reporter (Nern et al., 2011).

Last but not the least, pericyte degeneration associated with BBB dysfunction is found in a spectrum of CNS disorders, including AD and other dementia (Sweeney et al., 2016). Therefore, the identification of *Atp13a5* as a specific BBB pericyte marker will advance our studies in vascular contributions to AD and dementia. Beyond the application of the transgenic model, the identification of *Atp13a5* as a brain pericyte-specific marker can be used to develop other genetic tools, such as ATP13A5-specific antibodies, and new viral vector designs to target BBB pericytes more specifically with artificial promoters (Jüttner et al., 2019). These tools will have more broader applications, perhaps even in human clinical studies if *ATP13A5* can be validated as a human CNS pericyte marker (Noguchi et al., 2017).

## METHODS

### KEY RESOURCES TABLE

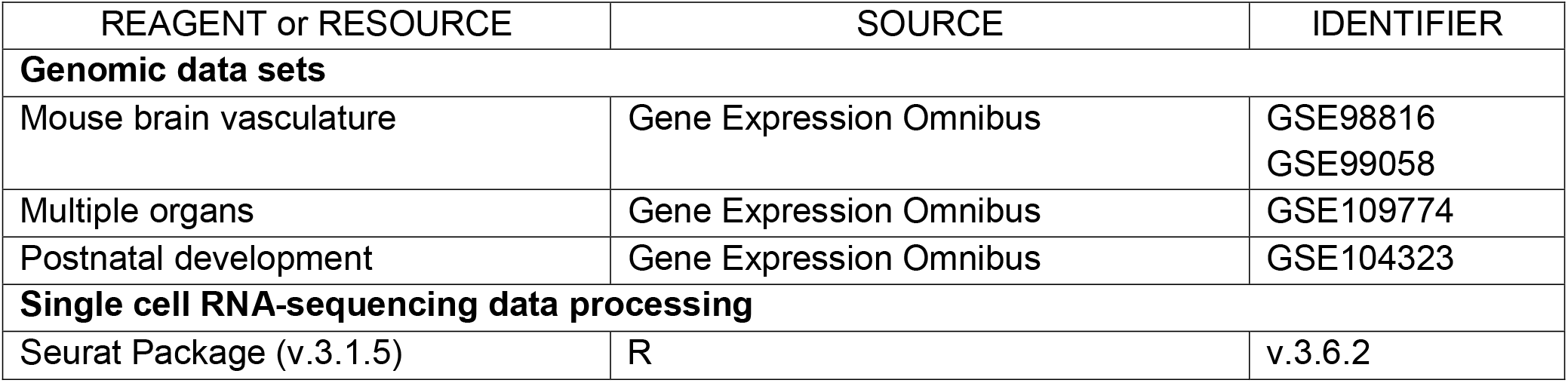
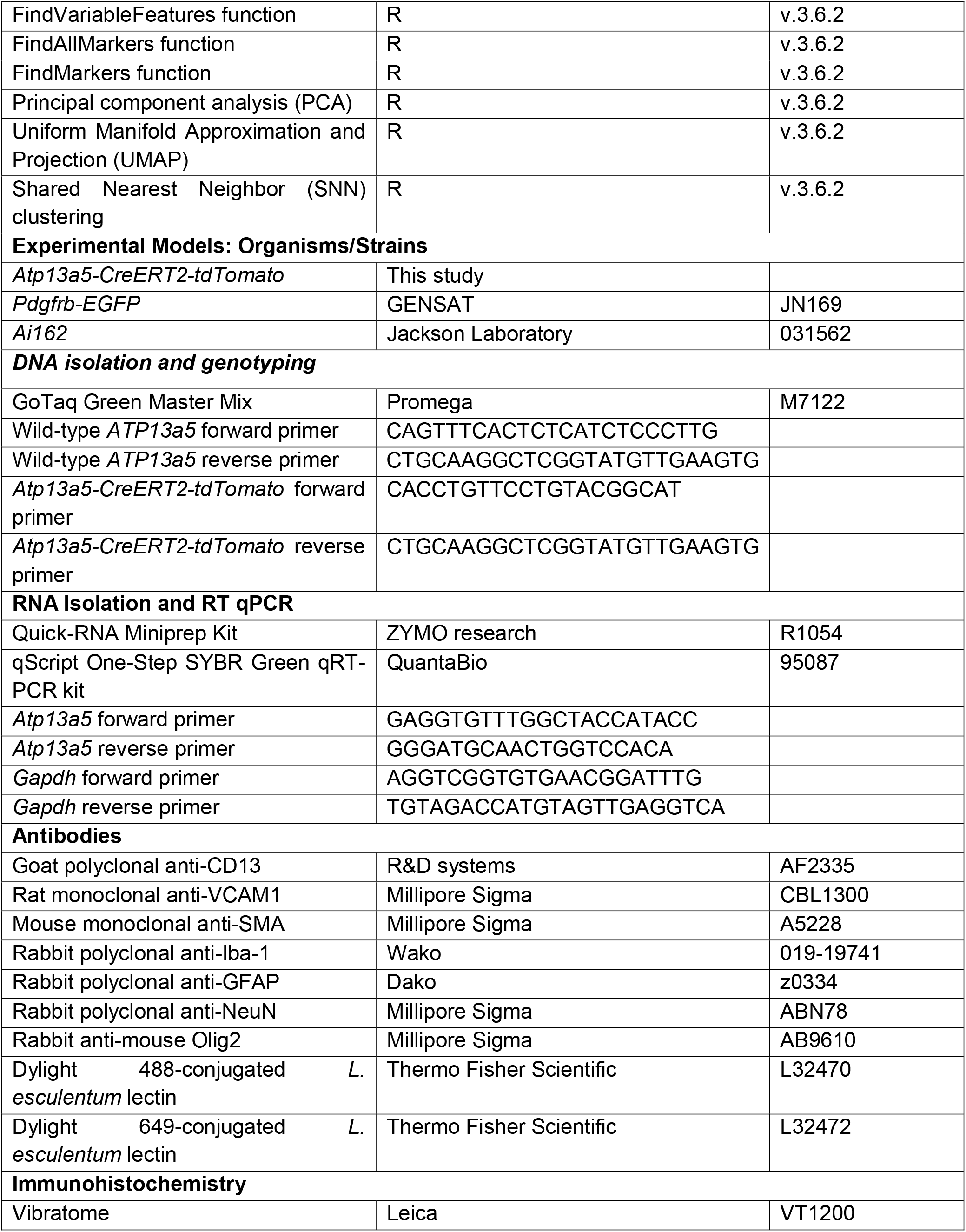
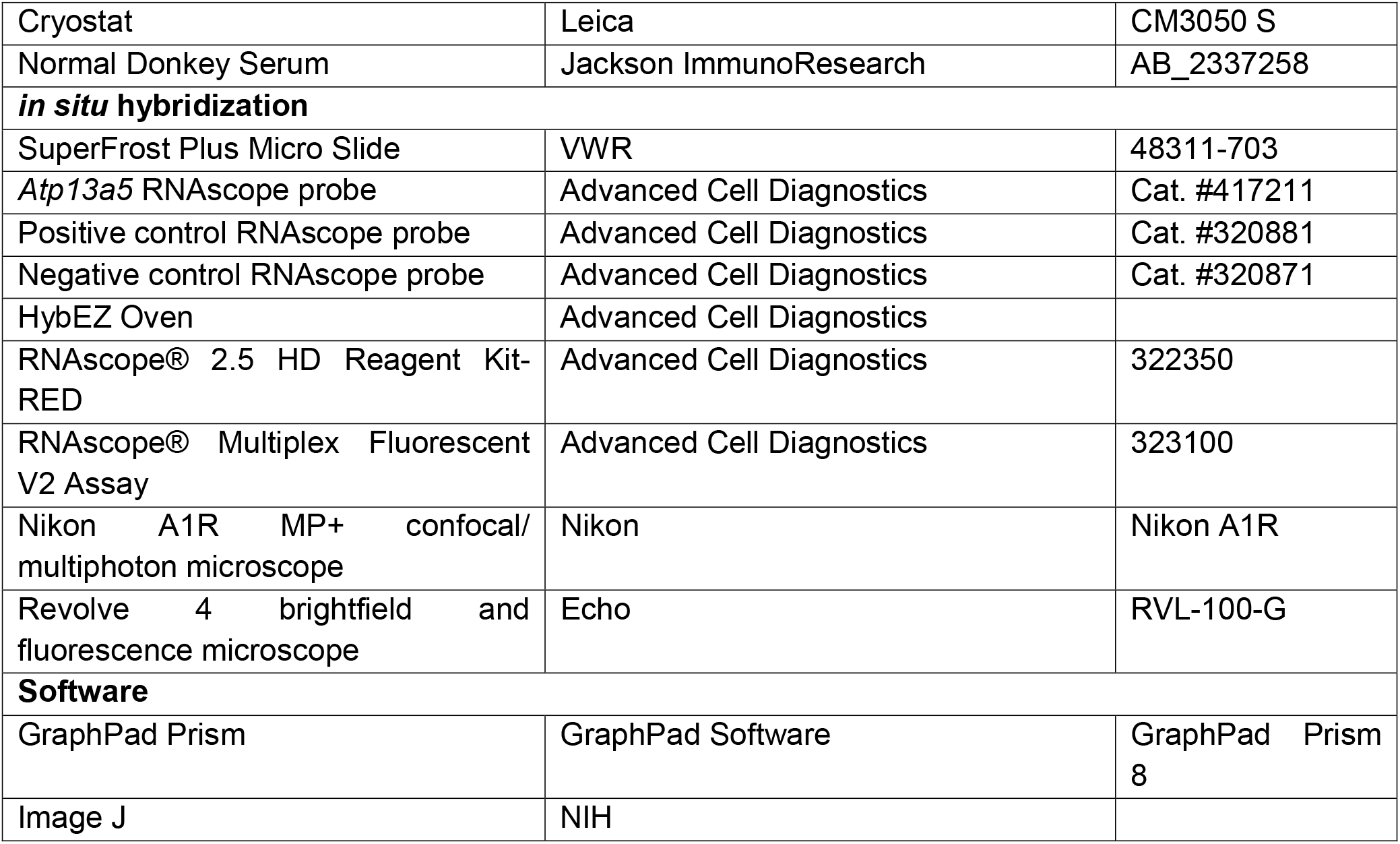

### CONTACT FOR REAGENT AND RESOURCE SHARING

Further information and request for resources and reagents should be directed to and will be fulfilled by Lead Contact Zhen Zhao.

### EXPERIMENTAL MODEL AND SUBJECT DETAILS

#### Animals

Mice were housed in plastic cages on a 12 h light/dark cycle with access to water ad libitum and a standard laboratory diet. All procedures were approved by the Institutional Animal Care and Use Committee at the University of Southern California and followed National Institutes of Health guidelines. All animals were included in the study. Male and female animals of 2–3 months of age were used in the experiments. All animals were randomized for their genotype information. All experiments were carried out blind: the operators responsible for the experimental procedures and data analysis were blinded and unaware of group allocation throughout the experiments. For all experiments, at least three independent mice were analyzed, which included both sexes and no apparent sex difference were observed.

##### Generation of the Atp13a5-2A-CreERT2-IRES-tdTomato knock-in model

To generate *Atp13a5-CreERT2-tdTomato knock-in mouse*, donor DNA templates encoding self-cleaving 2A peptide, CreERT2, internal ribosome entry site, tdTomato, and flp recombinase (*2A-CreERT2-Frt-IRES-tdTomato-Frt*) were synthesized. These sequences were flanked by 1184bp sequences and 1237bp sequences homologous to the last exon and 3’ UTR region of *Atp13a5* gene. In addition, the IRES-tdTomato sequence is further flanked by two flp recombinase target (frt) sites. Next, these donor vector containing the *2A-CreERT2-Frt-IRES-tdTomato-Frt* cassette, and gRNA (TTTTGGACTAGACTGTAACCAGG) were co-injected into fertilized C57BL/6N mouse eggs to generate targeted conditional knock-in offspring. The F0 founder animals were genotyped by PCR and sequence analysis, and 3 F1 mice were generated and further confirmed with southern blotting for both 5’ arm and 3’ arm insertion sequences (**Sup. Fig. 3A**).

Tamoxifen (Sigma, T-5648) administration in *Atp13a5-2A-CreERT2-IRES-tdTomato; Ai162* mice were performed at 40mg/kg per day for four consecutive days, as we described previously (Nikolakopoulou et al., 2019).

### METHOD DETAILS

#### Bioinformatics

##### ScRNA-seq data for mouse brain vasculature and multiple organs

For scRNA-seq dataset for mouse brain vasculature, we obtained the cell count matrix from Gene Expression Omnibus (GEO) with the series record GSE98816 and GSE99058 (Vanlandewijck et al., 2018). The data represent the expression levels of 18435 genes in 3186 cells. The mouse brain tissue was harvested for Smart-seq2 and sequencing was performed on a HiSeq2500 at the National Genomics Infrastructure (NGI), Science for Life Laboratory, Sweden, with single 50-bp reads (dual indexing reads).

For scRNA-seq dataset for multiple organs, we obtained the cell count matrix from GEO with the series record GSE109774 (The Tabula Muris Consortium et al., 2018). The data represent the expression levels of 23433 genes in 53760 cells. All organs were single-cell-sorted into plated using flurescence-activated cell sorting. Libraries were sequenced on the NovaSeq 6000 Sequencing System (Illumina) using 2 × 100-bp paired-end reads.

For scRNA-seq dataset for postnatal development, we obtained the cell count matrix from GEO with the series record GSE104323 (Hochgerner et al., 2018). The data represent the expression levels of 27933 genes in 24185 cells. The dentate gyrus from different ages was microdissected. All cDNA synthesis, library preparation, and sequencing were carried out as instructed by the manufacturer (10x Genomics Chromium Single Cell Kit Version 2). Libraries were sequenced on an Illumina HiSeq4000.

##### ScRNA-seq data preprocessing

The data processing of the scRNA-seq data were performed with the Seurat Package (v.3.1.5) in R (v.3.6.2) (Butler et al., 2018; Stuart et al., 2019). The basic scRNA-seq analysis was run using the pipeline provided by Seurat Tutorial (https://satijalab.org/seurat/v3.0/immune_alignment.html) as of June 24, 2019. In general, we set up the Seurat objects from different groups in experiments for normalizing the count data present in the assay. This achieves log-normalization of all datasets with a size factor of 10,000 transcript per cell. For different Seurat objects, FindVariableFeatures() function was used to identify outlier genes on a ‘mean variability plot’ for each object. The nFeatures parameter is 2000 as the default for the selection method called ‘vst’. These resulted genes serve to illustrate priority for further analysis.

##### Data processing

The dataset on all cells were used to scale and center the genes. First of all, principal component analysis (PCA) was used for linear dimensionality reduction with default computes the top 30 principal components. By applying the JackStraw() function, JackStrawPlot() function and ElbowPlot() function, we identified the principal components for further analysis. Then, PCA results were used as the input for the Uniform Manifold Approximation and Projection (UMAP) dimensional reduction.

We identified clusters of cells by a shared nearest neighbor (SNN) modularity optimization-based clustering algorithm. The algorithm first calculated k-nearest neighbors and computed the k-NN graph, and then optimizes the modularity function to determine clusters.

##### Determination of cell-type identity

To determine the cell type, we used FindAllMarkers() function with parameters min.pct and thresh.use set to 0.25 to find markers in each cluster and known marker genes that have been previously reported (Saunders et al., 2018) could be used to determine cell-type identity. These include, but are not limited to *Snap25* for Neuron, *Cldn10* for Astrocyte, *Mbp* for Oligodendrocyte, *Cldn5* for EC, *Kcnj8* for PC, *Acta2* for VSMC, *Ctss* for microglial, *Col1a1* for Fibroblast-like cell.

##### Cellular Biology Related Procedures

###### Fluorescence *in situ* hybridization

Fluorescence *in situ* hybridization was performed using the RNAscope technology (Advanced Cell Diagnostics, Hayward, CA). Tissue sample preparation and pretreatment were performed on fixed brains cut into 15 μm sections mounted onto SuperFrost Plus glass slides following the manufacturer’s protocol (ACD documents 323100). After dehydration and pretreatment, slides were subjected to RNAscope Multiplex Fluorescent Assay (ACD documents 323100). RNAscope probes for mouse *Atp13a5*, positive control and negative control were hybridized for 2h at 40°C in the HybEZ Oven and the remainder of the assay protocol was implements. Subsequently, the slides were subjected to immunohistochemistry. The fluorescent signal emanating from RNA probes and antibodies was visualized and captured using a Nikon AIR MP+ confocal/ multiphoton microscope (Nikon). All FISH images presented are projection of 10-image stacks (0.5 μm intervals) obtained from cerebral cortex, and a smoothing algorithm was applied during image post-processing (Nikon NIS-Elements Software).

###### Chromogenic *in situ* hybridization

Chromogenic *in situ* hybridization was performed using the RNAscope technology (Advanced Cell Diagnostics, Hayward, CA). Tissue sample preparation and pretreatment were performed on FFPE brain samples cut into 10 μm sections mounted onto SuperFrost Plus glass slides following the manufacturer’s protocol (ACD documents 322452). After deparaffinization and pretreatment, slides were subjected to RNAscope chromogenic ISH-Red Assay (ACD documents 322360). RNAscope probes for mouse *Atp13a5*, positive control and negative control were hybridized for 2 h at 40°C in the HybEZ Oven and the remainder of the assay protocol was implements. The *Atp13a5* Red signal was examined under a standard bright field microscope.

###### Immunohistochemistry

Animals were anesthetized, perfused and brains were removed and postfixed as we described previously (Nikolakopoulou et al., 2019). Brain, spinal cord, kidney, liver, and heart tissue were also collected, postfixed and cut at 35 μm thickness using a vibratome (Leica). After that, sections were blocked with 5% normal donkey serum (Vector Laboratories) and 0.1% Triton-X in 0.01M PBS and incubated with primary antibodies diluted in blocking solution overnight at 4°C. The primary antibody information is as following: Goat anti-mouse aminopeptidase N/ANPEP (CD13; R&D systems; AF2335; 1:100), Rat anti-mouse vascular adhesion molecule (VCAM1; MilliporeSigma; CBL1300; 1:200), Mouse anti-α-smooth muscle actin (SMA, MilliporeSigma; A5228, 1:200), Rabbit anti-mouse ionized calcium binding adaptor molecule 1 (Iba-1; Wako, 019-19741; 1:200), Rabbit anti-Glial Fibrillary Acidic Protein (GFAP; Dako, z0334; 1:500), Rabbit anti-mouse NeuN (Millipore, ABN78, 1:500), Rabbit anti-mouse Olig2 (Millipore; AB9610; 1:200). To visualize brain microvessels, sections were incubated with Dylight 488 or 647-conjugated *L. esculentum* lectin as we have described previously (Nikolakopoulou et al., 2019). After incubation with primary antibodies, sections were washed with PBS for three times and incubated with fluorophore-conjugated secondary antibodies. Sections were imaged with a Nikon AIR MP+ confocal/ multiphoton microscope (Nikon). Z-stack projections and pseudo-coloring were performed using Nikon NIS-Elements Software. Image post analysis was performed using ImageJ software.

##### Molecular Biology Related Procedures

###### DNA isolation and genotyping

Mouse genomic DNA was isolated from tail biopsies (2 - 5 mm) and following overnight digestion at 56□ °C into 100□μL of tail digestion buffer containing 10 mM Tris-HCl (pH 9.0), 50 mM KCl, 0.1% Triton X-100 and 0.4□mg/mL Proteinase K. Next, the tail will be incubated at 98 °C for 13 minutes to denature the Proteinase K. After centrifugation at 12000□rpm for 15 min, the supernatants were collected for PCR. Wild-type primers (432□bp): forward: 5’- CAGTTTCACTCTCATCTCCCTTG −3′; reverse: 5′-CTGCAAGGCTCGGTATGTTGAAGTG −3′. Knock-in primers (212□bp): forward: 5′-CACCTGTTCCTGTACGGCAT - 3′; reverse: 5′- CTGCAAGGCTCGGTATGTTGAAGTG-3′. The PCR conditions were as follows: 1) 94 °C for 3 min; 2) 35 cycles at 94□°C for 30 sec, 60□°C for 30 sec, and 72□°C for 35 sec; 3) 72□°C for 5□min. PCR products were separated on 2% agarose gel (**Sup. Fig. 3C**).

###### RNA isolation and real-time quantitative PCR

The mouse brains were harvested and frozen in dry ice and store at −0°C. Total RNA was isolated using Quick-RNA Miniprep Kit (ZYMO research, R1054) according to the manufacturer’s instructions and 10*μ*L of RNA was used for real-time quantitative PCR using qScript One-Step SYBR Green qRT-PCR kit (QuantaBio, 95087) according to the manufacturer’s instructions. *Gapdh* was used as an internal control for normalization. The primer information is as following: *Atp13a5* F’- GAGGTGTTTGGCTACCATACC; *Atp13a5* R’-GGGATGCAACTGGTCCACA; *Gapdh* F’ - AGGTCGGTGTGAACGGATTTG; *Gapdh* R’ - TGTAGACCATGTAGTTGAGGTCA. The PCR conditions were as following: 50°C for 5 min, 95°C for 30 s, and followed by 40 cycles at 95°C for 3 s and 60°C for 25 s.

###### Quantification and statistical analysis

Sample sizes were calculated using nQUERY, assuming a two-side alpha-level of 0.05, 80% power and homogeneous variances for the 2 samples to be compared, with the means and SEM for different parameters predicted from pilot study. All the data are presented as mean ± SEM as indicated in the figure legends and were analyzed by GraphPad Prism 8. For multiple comparisons, Bartlett’s test for equal variances was used to determine the variances between the multiple groups and one-way analysis of variance (ANOVA) followed by Tukey test was used to test statistical significance, using GraphPad Prism 8 software. A P value of less than 0.05 was considered statistically significant.

## Acknowledgments

The work of Z.Z. is supported by the National Institutes of Health (NIH) grant nos. R01AG061288, R01NS110687 and RF1NS122060, and BrightFocus Foundation grant no. A2019218S.

## Author contributions

XG and ZZ designed and performed experiments and analyzed data and wrote the manuscript. TG, SX, HW, MC, XX, BZ and FG performed experiments and/or analyzed data. JZ, JFC, DZ, AM and FG contributed to writing the manuscript. All authors read and approved the final manuscript.

