## Supplementary figures for "*Atp13a5* Marker Reveals Pericytes of The Central Nervous System in Mice"

**Supplementary Materials**

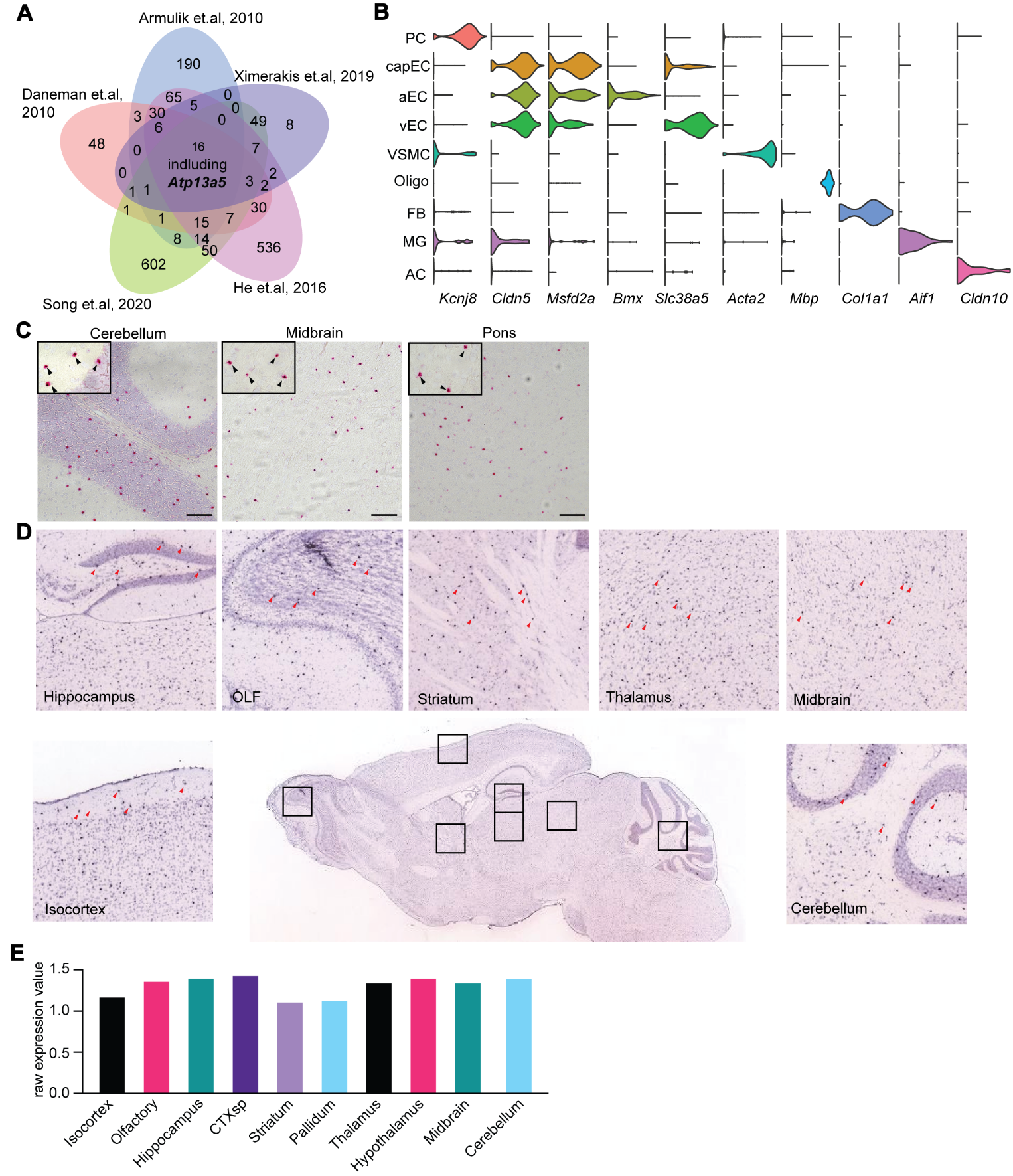

**Sup. Figure 1. *Atp13a5* expression in the mouse brain. (A**) Venn plot showing the overlaid genes between different datasets. **(B)** Violin plots showing gene markers that distinguish across vasculature cells. Genes are colored by cell types. PC: Pericytes; capEC: capillary endothelial cells; aEC: arterial endothelial cells; vEC: venous endothelial cells; VSMC: vascular smooth muscle cells; Oligo: Oligodendrocytes; FB: Fibroblast; MG: microglia; AC: astrocytes. **(C)** Representative images for *Atp13a5* mRNA expression (Red) in various mouse brain region. Scale bar: 100 µm. Sections: 10 µm thick. **(D)** ISH of *Atp13a5* mRNA expression in various mouse brain region from Allen Brain Atlas. (**E**) Raw expression value of *Atp13a5* mRNA expression in various mouse brain region. ISH expression data are from Allen Brain Atlas obtained from 56 days old adult male C57BL/6J mice (available from: <http://mouse.brain-map.org>). OLF, olfactory bulb; CTXsp, cortex subplate.

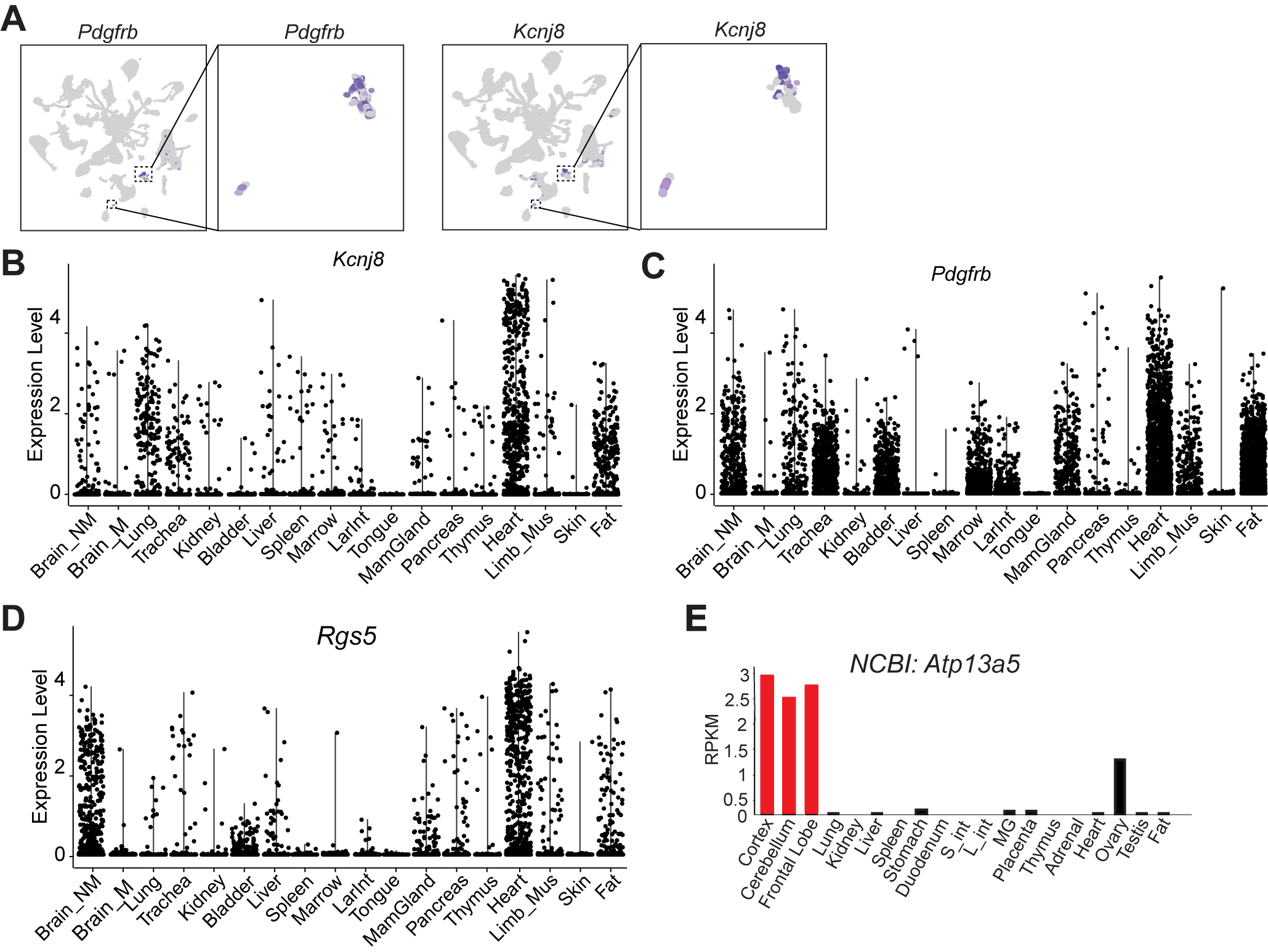

**Sup. Figure 2. Pericyte marker expression in different datasets. (A)** UMAP plots showing scRNA-seq data, colored by gene expression value, showing *Kcnj8* and *Pdgfrb* expression. (**B-D**) Violin plots showing *Kcnj8*, *Pdgfrb*, and *Rgs5* that distinguish across and within 18 tissues. Brain_M: Brain myeloid; Brain_NM: brain non-myeloid; LarInt: large intestine; Mus: muscle; MamGland: Mammary Gland. (**E**) Bar plot showing *Atp13a5* expression pattern in NCBI dataset. Red bar indicated the brain tissue. L_int: large intestine; S_int: small intestine; MG: mammary gland.

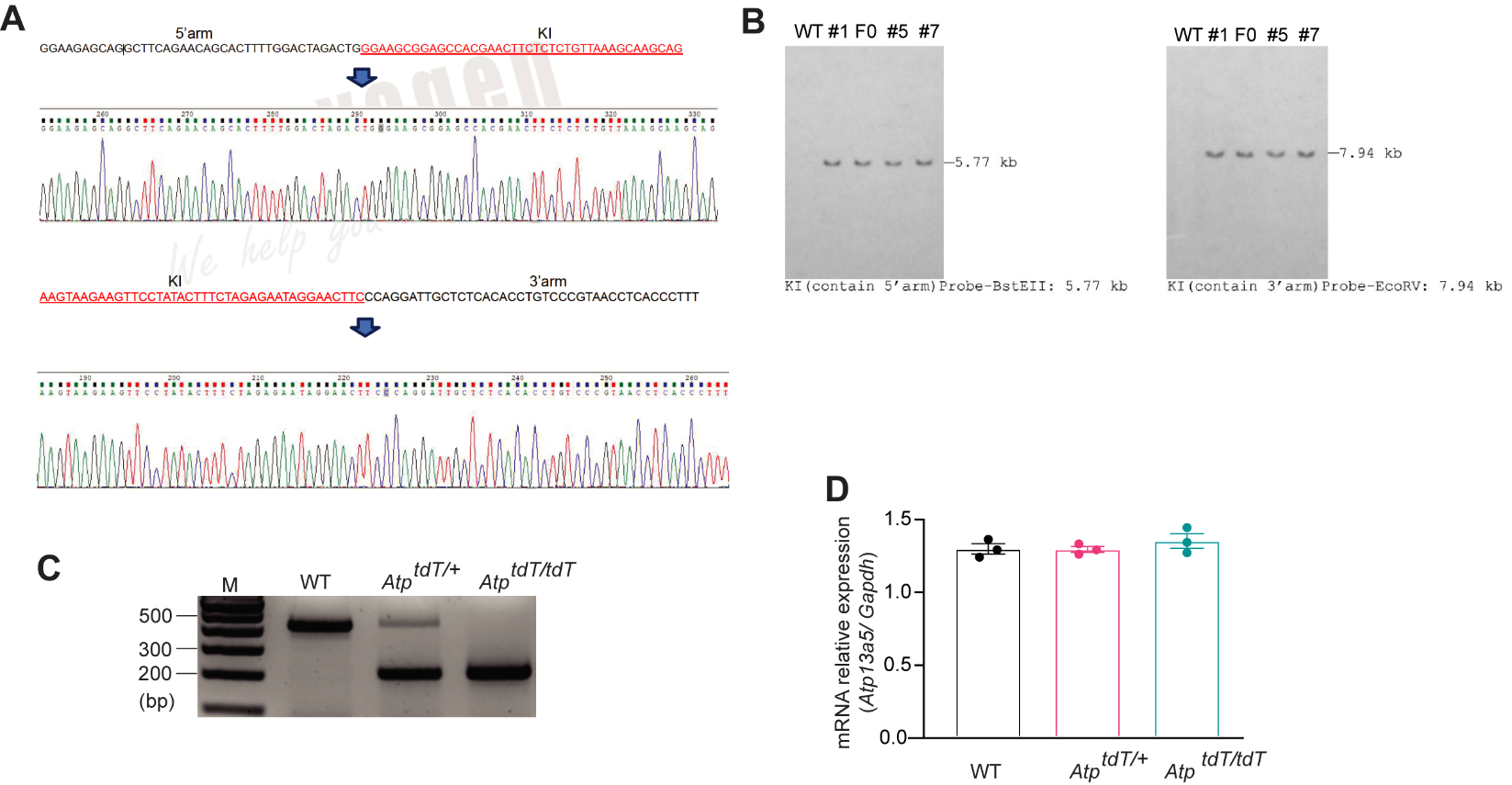

**Sup. Figure 3. Generation and validation of the *Atp13a5-2A-CreERT2-IRES-tdTomato* model. (A)** Sequencing analysis of F1 *Atp^tdT/^*^+^ mice showing the insertions at 5’arm (top) and 3’arm (bottom). No additional mutation or deletion were found. **(B)** Southern blotting analysis showing F0 and 3 F1 founders (#1, #5, #7) carrying the intact allele based on hybridization of probes targeting the 5’ and 3’ arms, compared to a WT littermate. **(C)** Genotyping result showing the genotype of WT (with a 432-bp band), *Atp^tdT/^*^+^ (with 432-bp and 212-bp bands) and *Atp^tdT/tdT^* (with a 212-bp band). *Atp, Atp13a5; tdT, tdTomato*. **(D)** The endogenous gene expression of *Atp13a5* relative to *Gapdh* in brains from 8-week-old WT, *Atp^tdT/+^* and *Atp^tdT/tdT^* mice (n = 3 mice each). Data are presented in mean ± SEM.

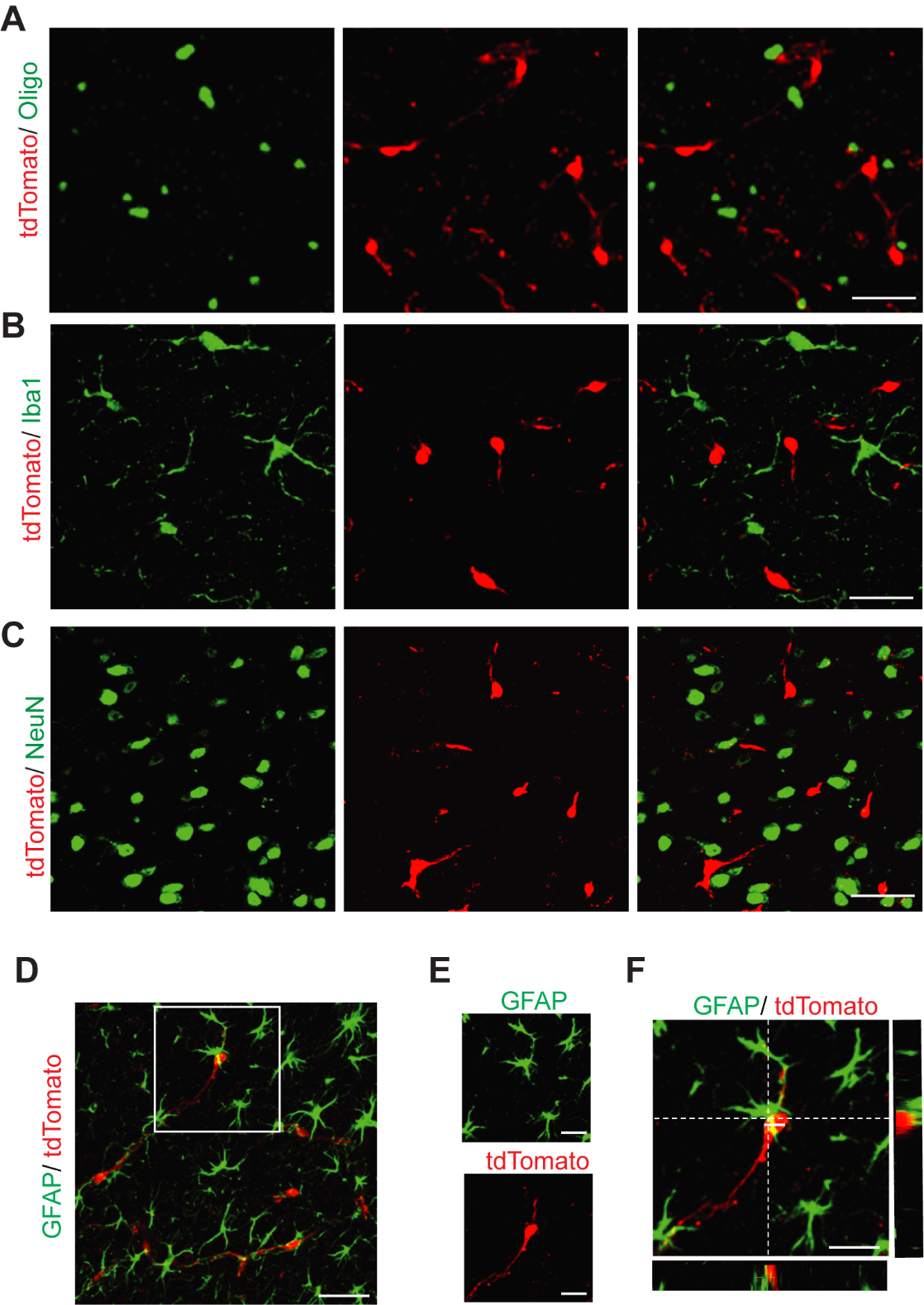

**Sup. Figure 4. Characterization of *Atp13a5-2A-CreERT2-IRES-tdTomato* mouse brain. (A-J)** Representative confocal images showing that tdTomato is not expressed in Oligo^+^ Oligodendrocytes (**A**), ionized calcium binding adaptor molecule 1 (Iba1)^+^ microglia (**B**), NeuN^+^ cortical neurons (**C**), and glial fibrillar acidic protein (GFAP)^+^ astrocytes (**D**). High magnification of boxed region in **D** is shown in **E**, and orthogonal view is shown in **F**. **A-D**: Scale bar: 50 µm. **E** and **F**: Scale bar: 25 µm. Sections: 35 µm thick.

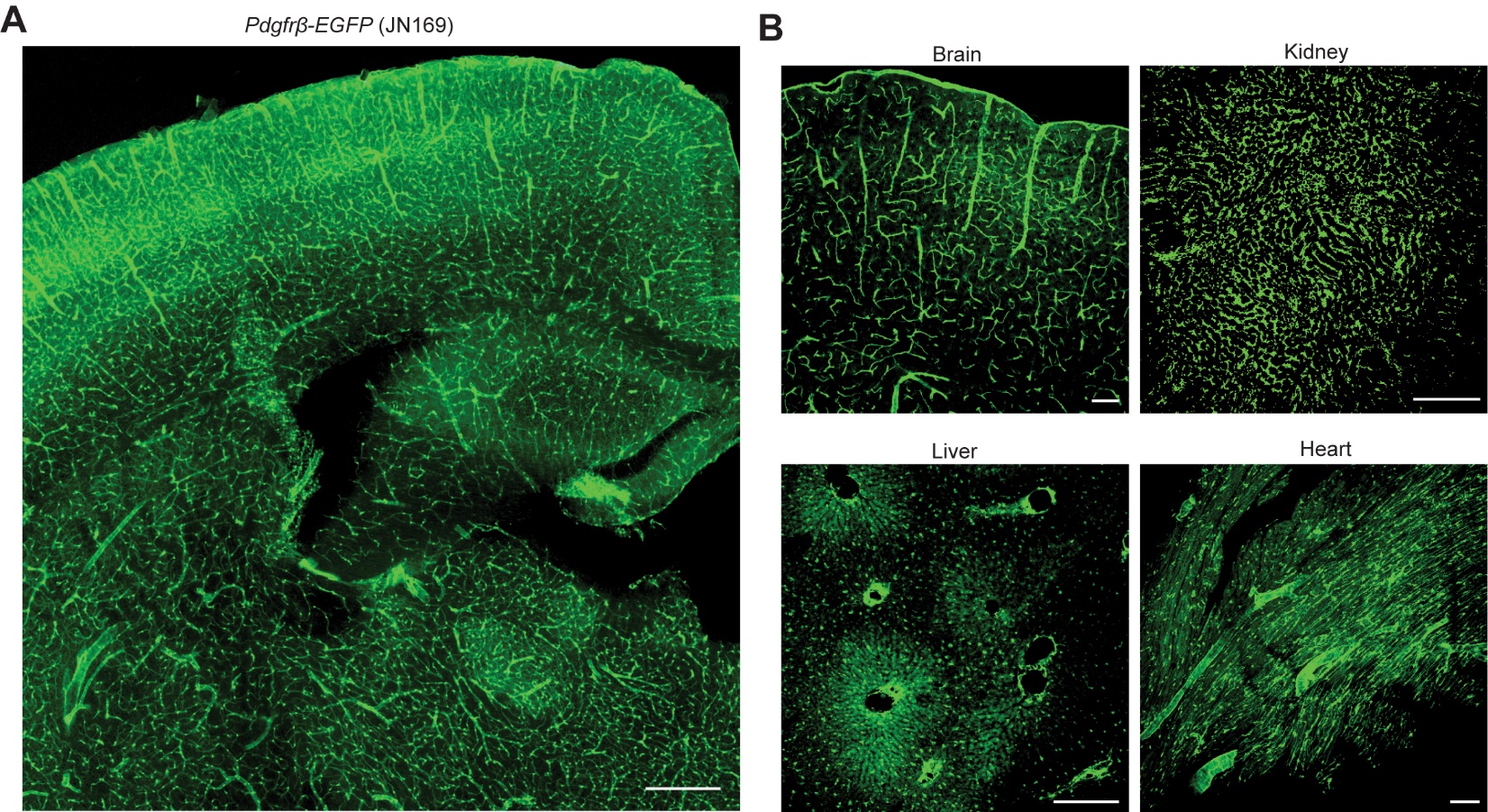

**Sup. Figure 5.** **EGFP expression in *Pdgfr***$\boldsymbol{\beta}$***-EGFP* mouse.** **(A)** A representative image of *Pdgfr*$\beta$*-EGFP* mouse brain. Scale bar: 200 µm. Sections: 35 µm thick. **(B)** EGFP reporter expression in brain (cortex) and peripheral tissues such as kidney, liver and heart. Scale bar: 100 µm. Sections: 35 µm thick.

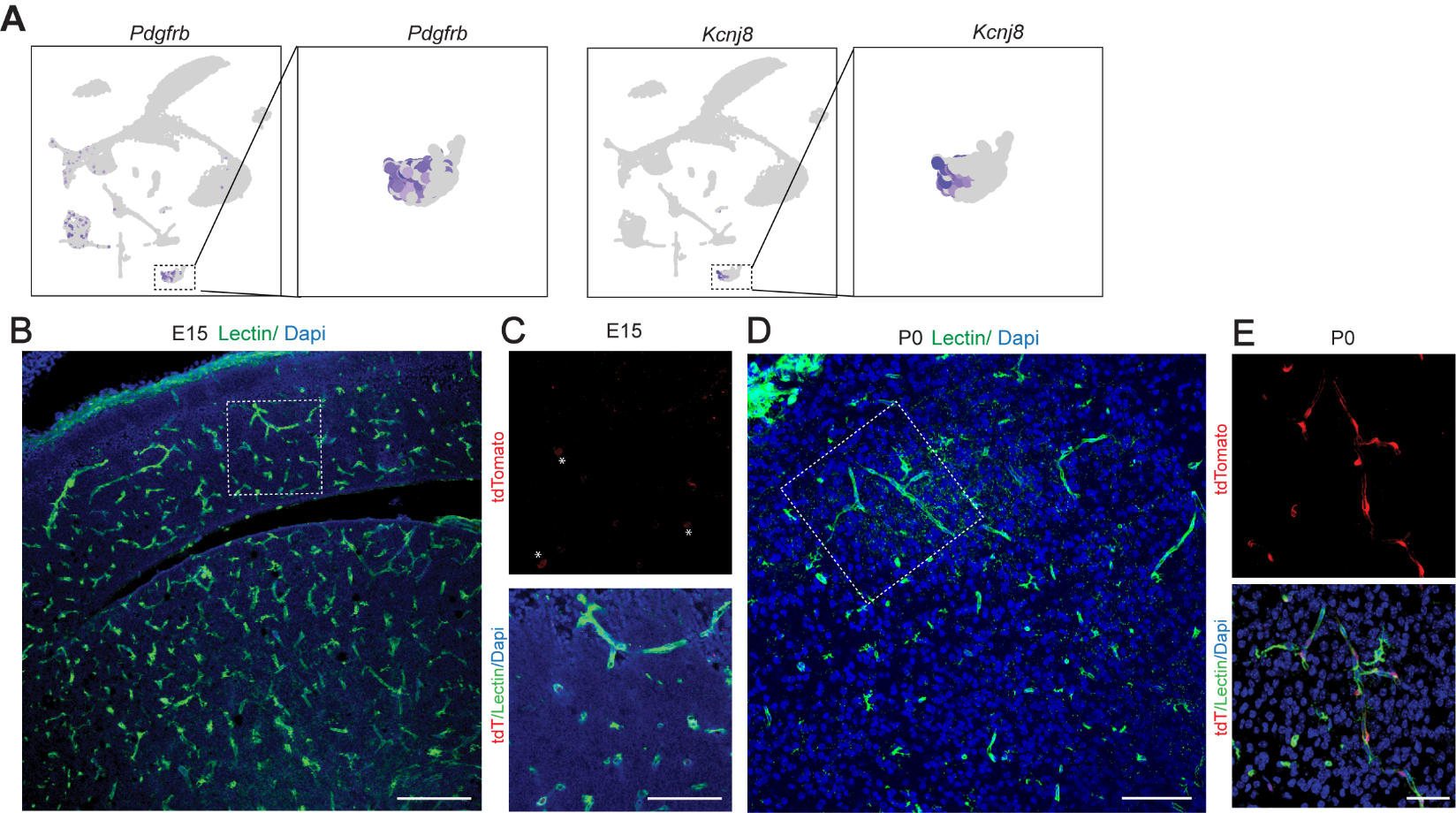

**Sup. Figure 6.** **Developmental regulation of *Atp13a5* driven tdTomato reporter expression.** (**A**) UMAP plots showing scRNA-seq data, colored by gene expression value, showing *Pdgfrb* and *Kcnj8* expression, based on dataset GSE95753. **(B)** A representative image of *Atp13a5-2A-CreERT2-IRES-tdTomato* mouse cortice at E15. Scale bar: 200 µm. Sections: 30 µm thick. **(C)** High magnification of boxed region in **B**. Asterisks: weak tdTomato signals detected in cortex at E15. Scale bar: 100 µm. **(D)** A representative image of *Atp13a5-2A-CreERT2-IRES-tdTomato* mice brain at P0. Scale bar: 100 µm. Sections: 35 µm thick. **(E)** High magnification of boxed region in **D**. Strong tdTomato signals can be detected in cortex at P0. Scale bar: 50 µm. Sections: 35 µm thick.
